# Disentangling the genomic complexity of the *Fragilariopsis cylindrus* (CCMP1102) genome

**DOI:** 10.1101/2022.07.14.500034

**Authors:** Kat Amy Hodgkinson, Jonathan Wright, Gonzalo Garcia Accinelli, Darren Heavens, Amanda Hopes, Thomas Mock, Cock van Oosterhout, Bernardo J. Clavijo

**Affiliations:** Earlham Institute, Norwich Research Park, Norwich, NR4 7UZ, UK; University of East Anglia, Norwich Research Park, Norwich, NR4 7TJ, UK

## Abstract

*Fragilariopsis cylindrus* CCMP1102 is characterised by a complex genome with significant levels of heterozygosity between haplotypes, > 35% repeats, and an unknown karyotype. This complexity hindered prior assemblies, which show coverage discrepancies indicative of incompleteness. Here, we use a k-mer spectra analysis to reveal the coverage signature for a third haplotype. We applied a novel haplotype-specific assembly method to reconstruct the *F. cylindrus* CCMP1102 genome, producing 10 fully assembled chromosomes capped by telomeres, and a putative chromosome with a single breakpoint. Our analysis shows triploidy, two cases of aneuploidy, and several truncations. We also present evidence that *F. cylindrus* reproduces sexually. Taken together, our analytical approach is capable of haplotype-resolved assemblies from structurally complex, polyploid genomes, making it suitable for complex genomes of non-model organisms, including those with unknown karyotype.

## Introduction

The sequencing of a wider range of organisms has been initiated to extend our knowledge of global biodiversity, exemplified by projects such as the Earth Biogenome Project (1), the 10KP project (2) and the 100 Diatoms project (https://jgi.doe.gov/csp-2021-100-diatom-genomes). These studies aim to understand the global diversity and evolutionary history of life and are expected to uncover many genomes with unprecedented complexity, ploidy variations and karyotypes. However, sequencing species with genomic structures that have not been assembled before, and for which no other genetic information is available, can be challenging. Genome assembly has typically produced a single pseudo-haploid reference consensus, collapsing haplotypes. Hence, it can miss important structural variations and lead to inaccurate gene annotations and artificial chimerisations that mislead biological interpretations (3). This is especially true for non-model organisms and genomes of specimens collected from the natural environment, which often contain high heterozygosity due to their outbred nature. Many such genomes also contain varying levels of ploidy, including aneuploidy, that may confound current genome assembly algorithms (4– 6). To address these challenges, haplotype-resolved *de novo* assemblies are required.

Diatoms are abundant in freshwater and marine environments. They are the most species-rich class of algae, contributing to about 20% global annual carbon fixation. *Fragilariopsis cylindrus* is an important indicator species of diatoms, native to polar oceans (7), with regular blooms in sea ice and open waters at the sea-ice edge. Over the recent years, it has become a model for the evolutionary adaptation of algae to polar oceans (8, 9). Its genome sequence became available in 2017 and was assembled *de novo* as a diploid by Sanger and PacBio sequencing. Its haplotypes were characterised by significant allelic divergence, however the assemblies were fragmented and contained uneven coverage, indicating that structural complexity may have been missed alongside some genomic content.

Recently, k-mer based analyses of reads have become more widely used to provide an indication of heterozygosity, repeat content, karyotype, coverage, and genome size, and read representation in assembled sequences (3, 10, 11). Reads are split into strings of k-length, and all distinct strings (k-mers) frequencies are counted. These read frequencies are compared to the counts of the distinct k-mers in the assembly. Integrating k-mer spectra into *de novo* assembly can avoid assumptions of ploidy and confirmation biases, providing an estimate of karyotypic characteristics for species with an unknown or varied karyotype. Additionally, the k-mer spectra can be used to assess and guide the assembly of haplotype-specific content (3).

We applied a novel assembly pipeline, the SDG-Threader, to the full karyotype re-assembly of *Fragilariopsis cylindrus*. CCMP1102 has a proposed 11 chromosomes, all of which are triploid but for two instances of aneuploidy. All chromosomes are capped by telomeres for at least one of the haplotypes. We also identified sites of recombination and several truncations, which end in telomeric sequence. The estimated genome size from this new assembly is 46Mbp based on haplotypes with no truncations. We show that this assembly pipeline is capable of assembling complex, non-model genomes with varied ploidy and can be applied to other diatoms for haplotype-specific assembly.

## Methods

### Genome sequencing

*Fragilariopsis cylindrus* (Grunow) Krieger CCMP1102 was obtained from the National Centre for Marine Algae and Microbiota (NCMA, East Boothbay, ME, USA), originally isolated from Southern Ocean seawater (64.08° S 48.7033° W, South Orkney Island Research Cruise, Station 12, 16th March 1979). Cell cultures were kept at +4 °C in f/2 medium. This culture was the same as used by Mock et al. (2017)(8) and Paajanen et al. (2017)(9). DNA was extracted using a cetyltrimethylammonium bromide (CTAB) based method modified from Friedl (1995)(12). For the paired end library a total of 2 µg of DNA was sheared targeting 1 kbp fragments on a Covaris-S2 (Covaris Brighton, UK), size selected on a Sage Science Blue Pippin 1.5% cassette (Sage Science, Beverly, USA) to remove DNA molecules <600bp, and amplification-free, paired end libraries constructed using the Kapa Biosciences Hyper Prep Kit (Roche, New Jersey, USA). This library was sequenced on an Illumina MiSeq. For long-read sequencing, DNA was not fragmented after extraction and was run on a High Pass Blue Pippin cassette targeting molecules >15kbp. This yielded approximately 2µg material for MinION library construction with an SQK-LSK109 kit. Half the total >600ng material was loaded onto a 106-REVD flowcell. All sequencing was produced at Earlham Institute (Norwich Research Park, Norwich, Norfolk, UK).

### Ploidy estimation

Initial 31-mer spectra-cn analyses with KAT (v2.3.4)(10) were performed between the Illumina data and the existing Sanger and PacBio assemblies to ascertain the existing level of resolution and review karyotype characteristics.

Smudgeplot (v 0.2.3)(13) was then used to corroborate ploidy level by analysis of the frequency relationship of k-mer pairs with a single nucleotide difference as a proxy for haplotype-level SNPs.

### Graph construction and long-read linkage

A canonical collapsed De Bruijn Graph (DBG) is first constructed in Sequence Distance Graph Threader (SDG-Threader)(14) from the short reads, which is then cleaned and simplified. Further repeat resolution is completed by finding unique paths between nodes representing haploid content, followed by canonical repeat resolution using long reads. Nodes longer than the 2Kbp are split into shorter nodes and every long read is then mapped to this length-split graph and represented as a thread. Nodes with 0.25<KCI<6 and less than 500 threads are selected, and every thread is used to create links between its consecutive selected nodes, creating the ReadThreadsGraph. From this, a number of heuristics create a collapsed graph, where, for two nodes to be adjacent, the support has to meet a criteria of minimum evidence. A final heuristic de-selects nodes that tangle haplotype paths. (See supplementary A and B for more details on how SDG was used to construct the assembly graph and perform the long-read linkage)

Telomeric sequences were identified by finding long reads anchored to nodes with no further onward links. Repeat motifs of ‘CCCTAA’ and ‘GGGTTA’, and their reverse compliments, were identified at both terminal ends of all chromosomes for at least one of the three haplotypes. These motifs were also identified at the terminal end of the truncated chromosomes.

### Manual analysis and verification of crossovers

We manually validated the potential cross-over points in SDG to ascertain whether they were mis-links or true cross-overs. We identified the supporting long-reads that threaded through nodes of differing K-mer Compression Index (KCI; see supplementary A), for which long-read linkage threaded 2x to 2x coverage nodes, 1x to 2x coverage and vice versa, in roughly equal quantities. We note that at these crossover sites, no 1x to 1x coverage linkages were present (See supplementary C). We used BANDAGE (v 0.8.1)(15) for graph visualisation and BLAST to map un-linked regions back to the assembly.

### Comparison with previous assemblies

KAT 31-mer spectra-cn plots were produced from the SDG and published assemblies to assess completeness and assembly composition. QUAST (v. 5.0.2)(16) was used to produce assembly stats and BUSCO (v. 5.3.2)(17) eukaryote_odb10 key genes aligned to assess completeness of the SDG assembly compared to the existing assemblies.

Local scale analysis of a structural difference was produced by isolating chromosomes 1 and 6 and aligning them to the published assemblies using NUCMER (v. 3.1)(18).

## Results

### Genome sequencing

The MiSeq sequencing run produced a total of 19,254,014 300bp paired reads, with an average fragment size of 500bp. The Nanopore run produced 746,568 reads, with an average size of 15.3Kbp and an N50 of 22.0Kbp.

### K-mer spectra analyses and ploidy estimation

The Smudgeplot analysis found 86% of the pairs show an AAB configuration, whereby A is present in two copies, indicating a triploid genome (Figure S1). A further 10% of pairs are in an AB configuration, where the k-mer pair does not have a third equivalent. These k-mers may be from a locus where only two copies exist, or the third copy may have more than 1 nucleotide difference and was therefore not tallied.

The 31-mer spectrum of the Illumina reads shows three distinct peaks, representing unique content (completely heterozygous), two copies (homo-/heterozygous) and three-copies (completely homozygous) (Figure S1). Unique genomic content appears at a fundamental frequency of 61. This is flanked by two further distributions at harmonic frequencies of 122 and 183 from the fundamental distribution, representing content twice and three times more frequent in the reads than the unique content.

K-mer spectra of the published assemblies shows some content is missing and some content is duplicated (Figure S2). This is reflected in the previous genome size estimate of 61.1Mb, which is possibly due to the chimerisation of haplotypes into one collapsed pseudo-haplotype.

### Assembly graph construction

The contig assembly contains 81Mbp in 6,541 contigs, with a largest contig of 308Kbp. The graph has an N50 of 29Kbp, N75 of 15Kbp and a GC content of 39%. K-mer spectra analysis confirmed all the genomic content was still present in the graph at this stage.

### Long-read-based haplotype reconstruction

Manual analysis of the linkages confirmed that the graph was mostly linearly structured as 2x coverage vs 1x coverage bubble sides. These indicated the presence of 2 identical haplotypes collapsed in one bubble side, and a heterozygous haplotype on the opposing side. These bubble sides sometimes terminate in a 3x coverage region, which indicates all three haplotypes are almost identical during these runs (Figure S3).

### Genome structure

The final graph contained 82Mbp, of which an estimated 46.6Mbp is haplotype A (Figure 1), which we suggest as the estimated genome size. This estimate is within a mutually complimentary range produced by qPCR of target genes (57.9Mb ± 16.9Mb; (8)). Homozygous content is collapsed but individual haplotypes can be retrieved from this graph. The graph contains 452 contigs, of which the largest is 2Mbp. N50 is 527Kbp and N75 318Kbp, with a GC content of 38% and 2,136 Ns per 100Kbp. This graph (Figure S4) is organised into 12 components, and its analysis enabled us to fully reconstruct the karyotype for *Fragilariopsis cylindrus* CCMP1102.

**Fig. 1.**
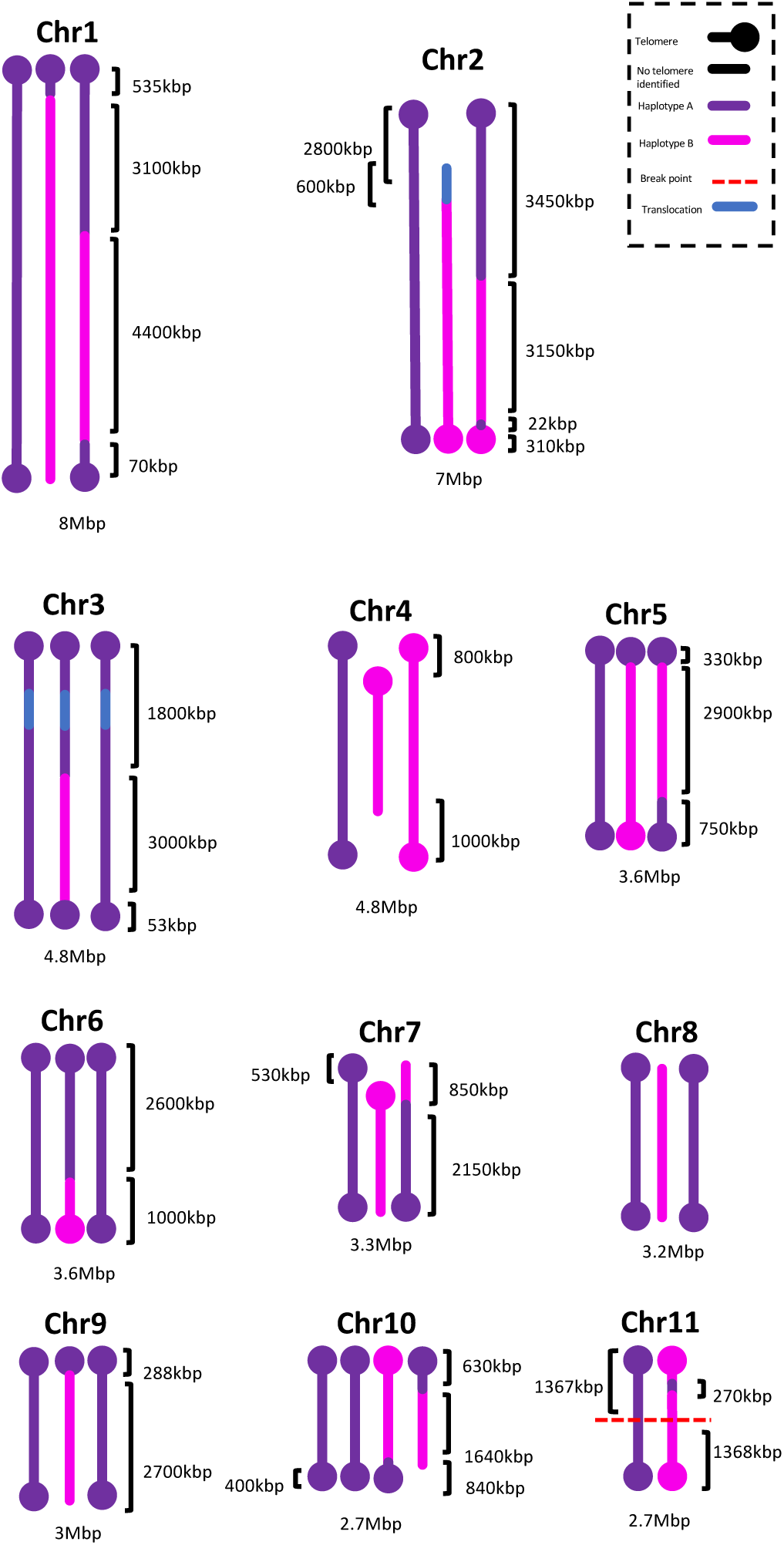
Proposed *Fragilariopsis cylindrus* karyotype. Chromosomes have been coloured based on haplotype identity. All content identical to haplotype A is coloured purple, while content heterozygous to A is coloured fuchsia. Where telomeres have been identified, the chromosome terminates in a blob, otherwise it remains open ended. Where Chr2 and Chr3 share a transposable element, this region is highlighted in blue. Chr11 has a break point where the two halves of both haplotypes remain unlinked. Estimated chromosome sizes and scales are shown for key features such as runs of homozygosity, crossovers and truncations. From left to right, chromosomes 1-9: A, B, X. Chromosome 10: A, A’, B, X. Chromosome 11: A, B.

The genome contains a proposed 11 chromosomes (Figure 1), capped by telomeres at both ends of all chromosomes for at least one haplotype, and telomeres at the truncations of chromosomes 4 and 8. Chromosome 10 is aneuploid with 4 haplotypes, while chromosome 11 contains 2 haplotypes. We note we could not establish a linkage between the two fragmented halves of the proposed chromosome 11, but we also could not detect telomeres on the unjoined ends, suggesting a potential linkage was missed.

A total of 5.6Mbp were detected as runs of homozygosity across 5 chromosomes, all of which occurred towards the telomere.

### Evidence of recombination

Recombination was identified on 5 chromosomes; chr1 has 2 crossovers, chr2 has 3 crossovers, chr5 has 1 crossover, chr8 has 1 crossover, and chr10 has 1 crossover.

### Validation and comparison to previously published versions

Initial k-mer spectra of the published assemblies versus the Illumina reads highlighted a proportion of missing heterozygous and 2x coverage content, which was more pronounced in the PacBio assembly (Figure S2). This totalled 11.8Mbp of heterozygous and 5.6Mbp of 2x content. This indicates that the goal of achieving a partially phased assembly was not met by the PacBio assembly, as large amounts of the target heterozygous content has been discarded. Only 4Mbp of heterozygous and 2.8Mbp 2x coverage was missing from the Sanger assembly, however some of this heterozygous content may be discarded contamination.

Our new assembly contains 10 complete and 1 fragmented chromosomes (Figure S4). Previous assemblies, were fragmented and contained no fully reconstructed chromosomes. Even when the purpose of our methods is not to provide fully ungapped reference sequences, the assembly contained 84% of the 255 key Eukaryotic BUSCO genes, similar to the Sanger assembly (84%) and higher than the PacBio assembly (81%).

Small scale local analysis of the published assemblies revealed some chimerisations between haplotypes and chromosomes (Figure S5). The collapsing of 3 haplotypes into a single pseudo-haploid reference sequence of an expected diploid genome may explain why the estimated genome size is 10Mb larger than our assembly.

## Discussion

### *Fragilariopsis cylindrus* karyotype

We present the first full haplotype-specific karyotype of the structurally complex diatom *F. cylindrus*. We discovered a third, previously unknown, chromosomal copy in addition to aneuploidy, and chromosomal truncations. While this could be an isolated case, it suggests diatom genomes complexity may have been systematically underestimated due to technical limitations in sequenceing and assembly methods.

We propose that this *F. cylindrus* strain (CCMP1102) is the result of meiotic non-disjunction between two sibling parent cells (Figure 2). Both parents likely originate from the same clonal lineage due to the low frequency of SNPs between syntenic regions, which were collapsed into runs of homozygosity in the assembly graph. One parent cell underwent normal meiosis, resulting in four 1n, recombined gametes as expected. However, the other parent cell underwent an aberrant meiosis I, likely due to failure of the checkpoint machinery, which resulted in non-disjunction and a 4n daughter cell going into prophase II. Meiosis I is a critical multi-step process resulting in genetic recombination and chromosomal reduction that precede gamete formation. Chiasmata formed at the sites of recombination play a role in disjunction mediation (see (19) and references therein), so recombination, or lack thereof, is closely associated with proper chromosomal disjunction. A failure to recombine is directly linked with aberrant miotic division (19, 20), and so these chromosomes remained in their parental structure. As a result, two 2n gamete cells were produced by the end of meiosis II.

**Fig. 2.**
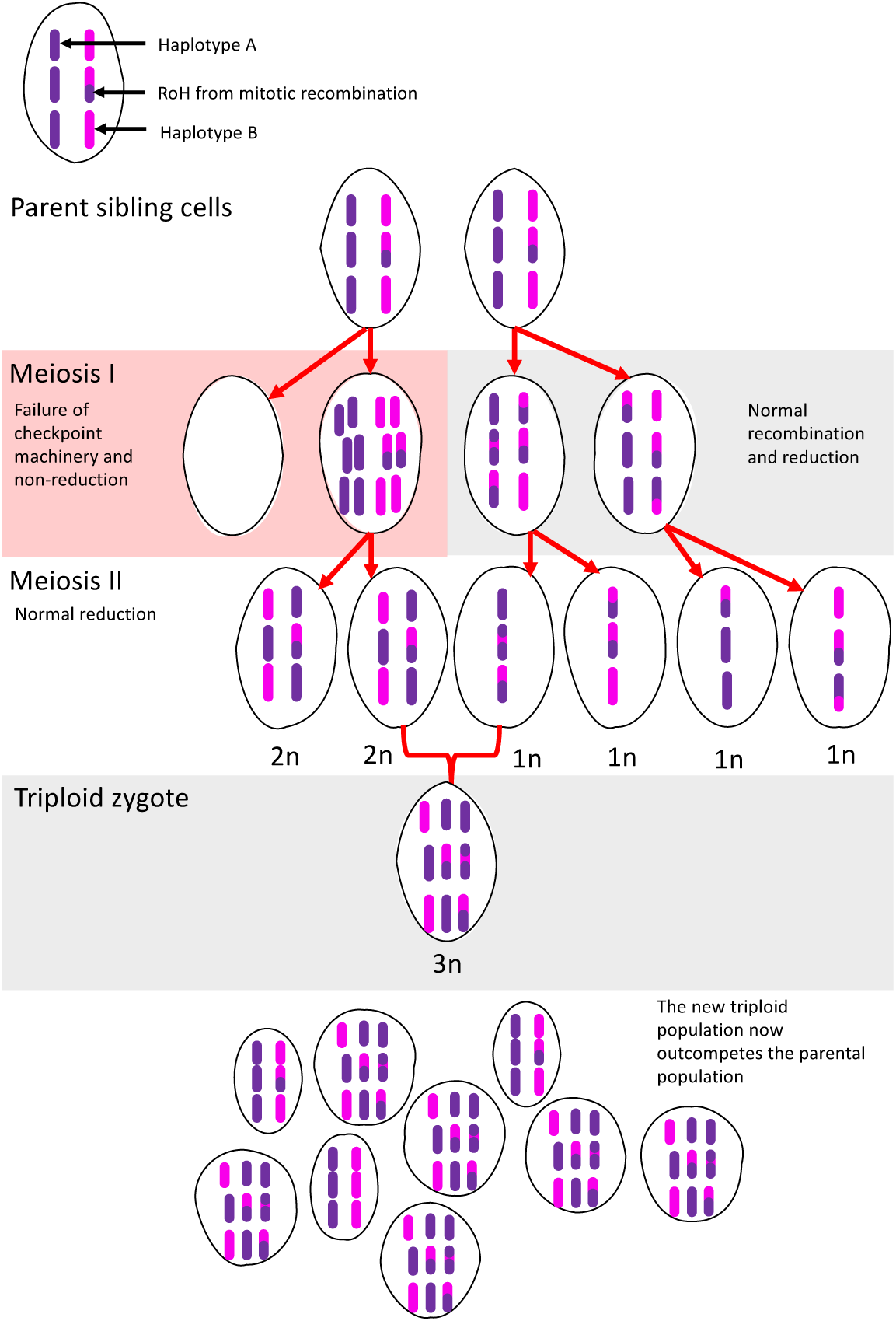
Non-reduction resulting from aberrant meiotic division. Aberrant meiotic division is represented on the left (red box), in contrast with the normal process on the right. The product of non-reduction during meiosis I are two daughter cells containing 0n and 4n, respectively. After meiosis II, this results in two 2n gametes, which upon fusion with a normal 1n gamete, produce a triploid. RoH = Run of Homozygosity resulting from mitotic recombination in the parental generation.

We note at the time of writing that this clonal individual has been in culture for over 40 years, and although we cannot conclude whether this individual is representative of natural populations, its genomic structure has been consistent throughout its sequencing history. Sexual recombination has not been observed in this species in laboratory cultures, and this may be in part due to the triploidy of this strain. Triploidy is an unstable state for a genome and causes problems with meiotic division that can result in sterility. Polyploid gametes formed from aberrant division during meiosis have been observed amongst diatoms (21–24), but the production of auxospores from a triploid have not been documented. However, recombination occurred at the expected frequency of 1-4 times on 5 chromosomes, presenting evidence that this species does sexually reproduce. Obligatory crossovers during Eukaryotic recombination are thought to ensure correct segregation during meiosis I and are expected at least once on every chromosome (25). As for the remaining chromosomes, we were unable to detect recombination either due to the location of the recombinant site amongst the homozygous runs, or because the inherited chromatid was non-recombined.

Otherwise highly differential haplotypes sometimes contained large expanses of homozygous content. Recombination rates generally increase from centromere to telomere (26), with linkage disequilibrium dissipating with distance from the centromere (27). This initially appears to contrast with our findings, where the homozygous regions were often positioned at the ends of chromosomes, capped by telomeres. However, it is consistent with non-crossovers during mitotic recombination of the parent cell, which results in loss of heterozygosity and fixation of alleles (28). In rapidly evolving organisms, non-core regions are associated with higher levels of mutation, whereas core regions are retained across haplotypes through mitotic recombination as runs of homozygosity (29). Future works may explore whether this is the case for *F. cylindrus*, and core genes, potentially linked with clonal reproduction, are located in these regions.

We can only speculate the origins of the aneuploidisations of 4n chromosome 10 and 2n chromosome 11. Although triploidy is found across many taxa (30–32), it is typically an unstable state and aneuploidy is often attributed to chromosomal instability caused by changes in the balanced stoichiometry of protein complexes (33). Due to the cell reverting to a euploid genome during clonal reproduction, it is possible a haplotype of chromosome 11 was lost. As chromosome 11 is the only diploid chromosome, future works may explore whether its genes are linked with sexual reproduction, which would give a reason as to why this euploidisation has occured.

Both aneuploidisation and truncation may also be attributed to non-disjunction. Truncated accessory chromosomes are shown to successfully inherit in model plants (34, 35), so it is possible for truncated chromosomes to be inherited through meiosis. Truncation occurs at broken chromosomal ends where repair machinery fails and instead, ends are capped with telomeres to maintain stability. Partial loss of genes from euploidisation or truncation likely have little effect on the survival of the cell as all core genes are still present in the necessary minimal dosage.

### Haplotype-specific assembly implications

The CCMP1102 assembly is a unique model consisting of two haplotypes with natural levels of variation, plus a third, recombined haplotype. This could have wide implications for studying the dosage effects of genes and adaptability. Investigating the dosage effect on the inversion of the haplotype, and whether the *cis-trans* regulation changes the function of genes from the evolved phenotype, can now be investigated using this assembly.

In previous work, a greater than fourfold significant unequal biallelic expression pattern was computed across 45% of diverged alleles when subjected to stress (8). This was interpreted as differential allelic expression (DAE) in the diploid assembly. However, along the length of most of the chromosomes we see three copies in two distinct haplotypes, each parent and a recombinant. Expression of alleles in polyploid plants often follows the dosage of haplotypic ratios (36), and polyploidy can increase the potential variation in expression levels (37). Therefore, firstly the dosage of allele expression of the polyploid must be established (36) before testing the effect of DAE across multiple conditions. More work needs to be done to understand the effect of the third haplotype on expression levels to be able to confirm or refute results from previous analyses (8).

A key aspect of assembly is the estimation of expected ploidy levels (38), but this may be difficult to predict for genomes assembled *de novo*. The k-mer spectrum provides a bias-free estimation method directly from raw sequencing reads, which enabled us to identify a third copy in the coverage distribution and apply a suitable assembly strategy. K-mer spectra can also quantify the completeness of assemblies by comparison to the raw sequencing read constituents (10). K-mer based methods have been shown to account for the complexities and levels of heterozygosity associated with wild genomes (39), and we have confirmed they are scalable to higher levels of ploidy.

Such complex, non-model *de novo* genome assembly requires copy number sensitivity, which the methods used for prior genome assembly did not have (8, 9). This forced diploidisation, which may have caused the chimerisation of the *F. cylindrus* haplotypes. The construction of a pseudohaploid reference consensus may also lead to loss of lower-coverage, unique content as haplotypes are collapsed into a consensus sequence.

High levels of heterozygosity complicate the assembly procedure, so whilst many assembly methods typically work well on model species with low heterozygosity, variation found in natural populations is lost. Conversely, our methods are more labour intensive and require more precise parameterisations, but when high coverage short reads or PacBio HiFi reads are combined with longer reads such as those from Oxford Nanopore Technologies (ONT), this recipe is capable of retaining all heterozygous content and resolving low-coverage or repeat-rich regions. This results in the chromosome scale, haplotype-specific assembly of non-model organisms.

### Conclusion

*F. cylindrus* CCMP1102 has a complex, triploid karyotype with high levels of heterozygosity, instances of aneuploidy and truncation. It reproduces sexually as is evident by the sites of recombination identified here. This genome was previously unable to be fully resolved due to technical limitations, causing important structures to be missed. These limitations are now being addressed in the genomics community through increasingly accurate, and longer scale, sequencing reads and methods of ploidy detection and haplotype-specific assembly. With the instigation of large-scale genome sequencing projects (1, 2), careful analysis of genome content followed by validation of karyotype structure will be key to correctly representing haplotypic diversity.

## Data and code availability

The raw reads generated for this project are deposited within BioProject PRJEB51975.

## Author Contributions

The study was designed by KAH and BJC. AH produced the cultures. DH performed DNA extraction and Nanopore sequencing. KAH, JW, GGA and BJC analysed sequencing results, assembled the sequences, and performed all analyses on the assemblies. KAH and BJC produced the initial manuscript draft. CvO, TM and AH provided discussion and contributed to result interpretation. All authors contributed to the writing and approval of the final manuscript.

## ACKNOWLEDGEMENTS

This work was strategically funded by the BBSRC Core Strategic Programme Grant [BBS/E/T/000PR9818] Work by GGA and BJC was also partially funded by the BBSRC grant “OctoSeq: Sequencing the octoploid strawberry” [BB/N009819/1].

KAH acknowledges Natural Environment Research Council funded Aries DTP (NE/L002582/1). TM and CvO acknowledge the Natural Environment Research Council (grant nos. NE/K004530/1 and NE/R000883/1), The Leverhulme Trust (RPG-2017-364), and partial funding from the School of Environmental Sciences at the University of East Anglia, Norwich Research Park, United Kingdom.

