## Supplementary Material for "Disentangling the genomic complexity of the *Fragilariopsis cylindrus* (CCMP1102) genome"

### Supplementary Note 1: Assembly methods

#### A. SDG contig assembly.

The K-mer Compression Index (KCI) is an integral deterministic metric used by SDG. It is used to determine the copy number of a node as it would appear in the final resolved assembly, such that a KCI of 1 would be a unique node, 2 would contain collapsed duplicated sequence, 3 contains collapsed triplicated sequence, and so on. This can be used to identify heterozygous or homozygous nodes. KCI is calculated based on the ratio between the median coverage of a node and its fundamental k-mers (k-mers that only appear once in the graph), and a unique coverage that is typically set by the user as the peak of the unique content distribution of the reads (Figure S1). The KCI parameterisation in *F. cylindrus* is adjusted to account for triploidy, so that KCI = 1 is unique, or heterozygous sequence, KCI = 2 indicates collapsed duplicated sequence that is homozygous across two haplotypes, and KCI = 3 indicates triplicated collapsed sequence, or homozygous across three haplotypes.
Contig assembly is performed as described in the SDG-Threader walkthrough (https://github.com/bioinfologics/sdg-threader/blob/main/doc/tutorial.md).

```
01-dbg.py -o Fcyl_triploid31-b 8 --kci_k 63 -o Fcyl -p PE.prseq

02-clean.py -o Fcyl_triploid31 -u 50

03-short_repeats.py -o Fcyl_triploid31 -u 50

04-strider.py -o Fcyl_triploid31 -u 50

05-long_repeats.py -o Fcyl_triploid31 -l LR.loseq -u 50
--lr_min_support 3 --lr_snr 5 --lr_max_noise 5 --final_remap true
```

SDG's DBG module constructs a canonical collapsed DBG, which is then cleaned and simplified. The basic theoretical principle is that given a complete and error-free graph, every linear sequence in the genome should be traceable as a path through the graph, and the full assembly should pass through every node at least once. There are three major structures that the algorithms work to resolve: tips, canonical repeats and bubble sides. Linkage errors cause the formation of tips, dead end nodes that have only one input link from the previous node. Canonical repeats are repetitive nodes with a minimum of 2 input and output links, where the number of inputs and outputs is the same. Bubble sides are a pair of parallel nodes connecting to the same input and output nodes, forming distinctive bubbles in the graph.

#### B. SDG ReadThreads graph construction.

Until this point, the assembly graph should contain all the genomic content including collapsed repeat regions, where haplotypes are not easily separated due to their close identity. This assembly graph has increased contiguity but many paths that could give rise to mis-assemblies remain and chromosomes and haplotypes are entwined. The construction of a ReadThreadsGraph aims to make the resolution into a full karyotype possible by threading long reads through resolved sections of the graph.
In the first step, nodes longer than the set parameter size (default 2Kbp) are split into shorter nodes. This avoids incorporation of structural misrepresentations from the long nodes into the graph by mis-joins. Every long read is then mapped to this length-split graph and represented as a thread (i.e. a series of best-mapping nodes). These threads represent partial proposed solutions to the path-finding problem of haplotype reconstruction from a graph with collapsed repeats. To create a ReadThreadsGraph, a set of nodes is selected (by default nodes with 0.5<KCI<1.5 and less than 1000 threads) and then every thread is used to create links between its consecutive selected nodes. The ReadThreadsGraph then contains as many links between two nodes as threads where the nodes appear next to each other. Therefore from this, a number of heuristics create a collapsed graph, where, for two nodes to be adjacent, the support has to meet a criteria of minimum evidence. A final heuristic de-selects nodes that tangle haplotype paths. These are tips or repeat nodes that, if deselected, will allow their connected non-repeat neighbour nodes to become linearly connected.
Further selection or deselection of nodes can be manually introduced to finalise graph resolution. To aid in the resolution of some complex regions, deselection of specific links between nodes is also supported.

```
06-split_and_map.py -i Fcyl_05_long_repeats.sdgws -o Fcyl -u 50 -s 2000

07-thread_and_scaff.py -o Fcyl -u 50 -s 800 --min_kci .25 --max_kci 6
--min_hits 2 --min_links 5 --max_thread_count 500
--include_nodes 300:10000:0.25:2.5 --include_nodes 800:10000:0.25:6
--min_bp 130 --rrtg_edits rrtg_edits
```

From this point, if the graph still contains mis-joins, the graph can be manually viewed and manipulated to remove dubious linkages, repeat nodes or to reintroduce lost linkages and nodes. For this, two kinds of graph are used: the contig graph contains all information from the reads including every node, linkage, and repeat. This graph maintains all the linkages potentially lost in the scaffolded graph due to parameterisations, so can be used to re-link lost structure. The scaffolded graph only contains nodes that meet specific parameter requirements, including length, uniqueness and linkages, and as such, does not contain the full sequence. From this graph, when combined with the KCI, we can explore the full structure of the chromosomes, including size, karyotype, haplotypes, crossovers and telomeric positions. Deselected nodes from repeat regions are still present within the contig graph, so can be manually re-selected to reintroduce lost linkages. To manually analyse areas of the graph, we use a combination of the scaffolded graph to understand the overall structure, and the original graph to see the underlying sequences and all its linkages. This combination allows us to manually adjust the parametrisation to increase contiguity or understand complex structural features.

Sometimes, particular areas of the genome that are not well sequenced may have low coverage and fall outside the expected parameterisation, thus needing manual analysis and resolution. In other cases, mis-links may be caused by unresolved canonical repeats, which can be manually resolved by excluding repeat nodes or breaking links with low long-read support (Figure S6). This can resolve false links between chromosomes. Other mis-assemblies, such as fragmentation, can be resolved by aligning the fragmented regions back to the graph and tracing the long-read support through the original graph. Manual analysis also gives an idea of how the coverages are distributed across the assembly. Nodes with a KCI of 1 are unique and so heterozygous, and nodes with a KCI of 2 are collapsed duplicated sequence, or homozygous.

Manual analysis of the linkages confirmed that the graph was mostly linearly structured as 2x coverage vs 1x coverage bubble sides. These indicated the presence of 2 identical haplotypes collapsed in one bubble side, and a heterozygous haplotype on the opposing side (Figure S3). These bubble sides sometimes terminate in a 3x coverage region, which indicates all are almost identical during these runs.

We manually identified nodes that were not included in the scaffolded version, such as nodes that fell outside of KCI or length thresholds, and adjusted the parameterisation accordingly. Many of these nodes were not being included because they fell outside the KCI thresholds, which were originally set to be strict (KCI1: 0.8-1.2, KCI2: 1.8-2.2, KCI3: 2.8-3.2) to reduce noise in the graph. Some manual resolution of gaps was also possible by traversing repeat regions and selecting the contiguous path through long-read supported linkages. This gap resolution within the triple regions added in divergent nodes containing SNPs, which could later be used for SNP calling.

The final step uses the join_graph script to join the graph back together. This replaces some of the missing content so that the only missing nodes are those within unresolved repeats. Each haplotype can be outputted as a linear path traced through every node of the chromosome, including the shared homozygous nodes.

#### C. ReadThreads graph analysis.

We found 3 types of potential crossover in the scaffold graph that needed validating manually (Figure S7). These crossovers could have been potential errors in the algorithm causing problems with haplotype distinction, or may have been genuine crossovers between haplotypes that needed confirmation.

The first type of crossover occurred at the point that 2x and 1x nodes collapsed into a 3x node or short region. The long read support for these areas indicate linkages between 1x to 1x and 2x to 2x coverages only. No linkages between coverages were present, suggesting there are no haplotype crossovers. These areas are more frequent than the true haplotype crossovers and appear to show small stretches of sequence that are identical between all 3 haplotypes.

On most chromosomes we found between 1-4 crossovers where the double coverage became single coverage and vice versa. We investigated why this was occurring by finding the long-read support for the linking nodes. These long-read linkages strongly supported the linkages in the graph, with no linkages occurring between two 1x coverage nodes. This implied that one of the haploid genomes was switching between haplotypes, indicative of recombination events.

Other crossovers that were not confirmed as genuine contained high numbers of linkages between 1x coverage nodes, which were half that of the 2x to 2x linkages, as expected. Additionally, the 1x to 2x linkages were supported by comparatively fewer long reads. This kind of crossover was thus judged to be an algorithmic mislinkage.

In some regions of the graph, coverage changes suggested one copy was being lost. These changes occurred as 2x vs 1x to 1x vs 1x coverage change, or entire chromosomes that were missing a third copy. We hypothesised two potential scenarios for this; 1) the missing component was highly diverged and so the linkage was dropped, but should still be present in the graph; 2) the missing component was not present in the genome, implying a biological reason for the missing haplotype.

We tested the first hypothesis by aligning aneuploid components back to the graph to search for the missing third copy using the BLAST feature of Bandage. This resulted in no additional alignments in either the scaffolded or contig graphs. We then checked the k-mer spectra comparison of the unique components to see if there was content missing from the graph that was present in the reads, but all content was present. We concluded that the missing fragment of chromosome must be missing from the reads. This suggests a level of aneuploidy across some of the chromosomes, whereby some chromosomes are diploid and others are missing regions of the triploid chromosome.

We also found one chromosome with a signature of tetraploidy. On this chromosome we could see a triple-coverage scaffold diverge into the expected double-coverage and unique bubble sides (Figure S3). However, the unique bubble side reaches a complex repeat region, whereby it changes to a double-coverage bubble side. From this point, the chromosome contains four copies, two per bubble side. At the opposing end of the chromosome, the double-coverage bubble sides diverge into one triple-coverage and one unique region. As with the other aneuploid regions, we searched for alignments elsewhere in the graph for unlinked segments but found none. We also aligned the 1x coverage region to the whole graph to determine whether this 1x coverage segment aligned to another 2x coverage segment in the graph. However, no further alignments were identified and the 1x segment aligned only to the linked 3x segment.

We then traced the linkages through the aneuploid regions using the contig graph and the SDG long-read linkage support. From this we found 63 links supporting double coverage to triple coverage and 29 links supporting one of the double-coverage bubble sides to the single-coverage region (Figure S3). No links supported any other potential paths. This is consistent with three of the haplotypes converging into a homozygous region and one of the haplotypes diverging into a heterozygous region, as we see in the graph.

Our hypothesis is that this chromosome is showing a complex tetraploid aneuploidy; two of the haplotypes are full copies; the other two are alternate haplotypes with one of these being a recombined haplotype and one having lost a segment (Figure 1). However, which haplotype is recombined, and which is missing a segment, or whether they are the same haplotype, we cannot ascertain from the present data.

In two cases we found telomeric motifs at the ends of shortened haplotypes. We aligned the sequence preceding the telomere back to the full-length haplotype to find whether the segment had been truncated or whether there was a deletion. There was no alignment to the end of the full haplotype, suggesting a truncation of the shortened haplotype. We searched for unlinked regions using BLAST but no unlinked regions were found.

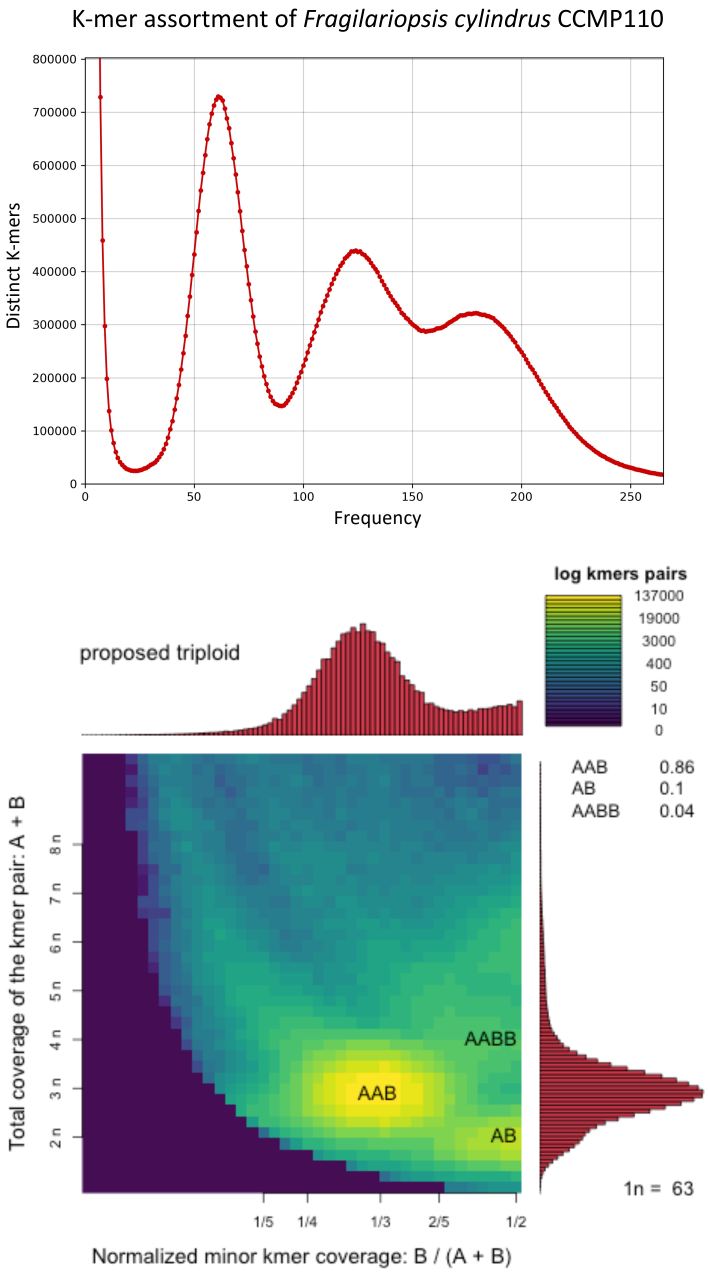


**Fig. S1.** *Fragilariopsis cylindrus* was found to have a triploid signature based on the assortment of its genomic k-mers. (Above) K-mer spectra analysis shows three distinct peaks, with the first representing heterozygous content and the two following peaks at harmonic frequencies from the first. (Below) Smudgeplot showing k-mer pairs with the proportion of AAB k-mer pairs is strongly indicative of triploidy.

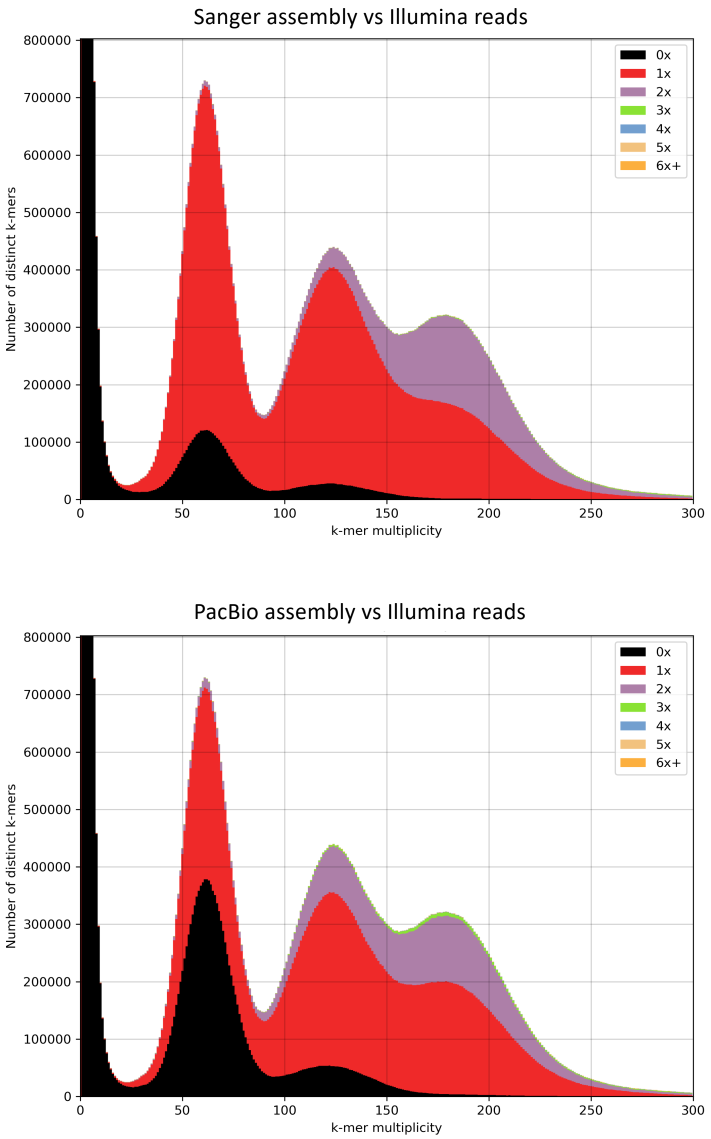


**Fig. S2.** The 27-mer spectras of the published Sanger and PacBio *Fragilariopsis cylindrus* assemblies versus Illumina reads. Black content from the first peak at a frequency of 62 represents unique, heterozygous content that is not included in the assemblies. This is more pronounced in the PacBio assembly. Some content is also missing from the 2x coverage peak at a frequency of 122. Duplicated sequence can be seen in purple and green.

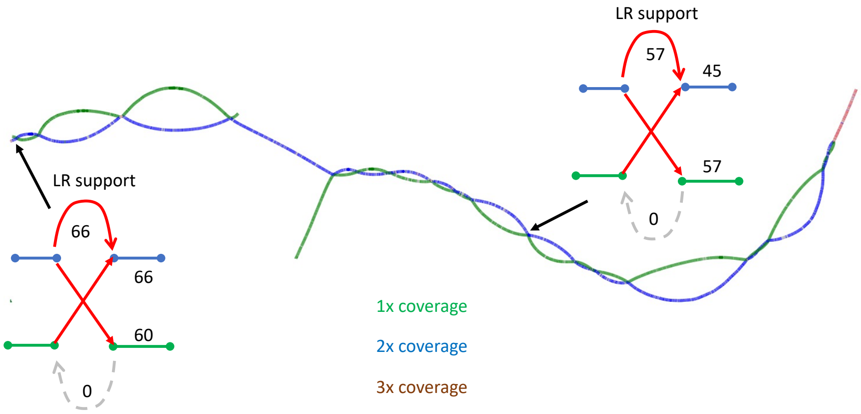


**Fig. S3.** Chromosome 2 crossovers. In the case that there is a recombination event of the haplotype, we would expect even coverage of the long-reads supporting those linkages. The two crossovers on chromosome 2 from 1x to 2x coverage show high amounts of long-read support. Similarly links between 2x to 2x nodes also show high levels of long-read support. However, no long-reads link 1x to 1x nodes.

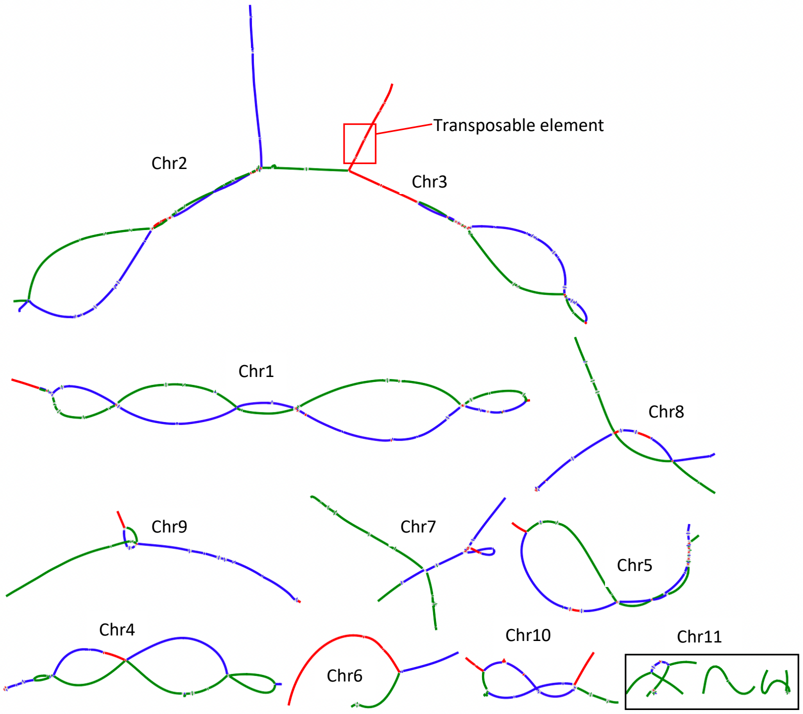


**Fig. S4.** Final Fragilariopsis cylindrus CCMP1102 assembly graph. Green represents heterozygous content, blue has 2x coverage and so represents 2 collapsed haplotypes, and red represents 3x coverage or over and so 3 or more collapsed haplotypes. Bubbles are formed where the haplotypes converge into a small shared node. The graph contains a confirmed 10 chromosomes and putative chromosome 11 with at least one haplotype capped at both ends by telomeres. A potential linkage between arms of chromosome 11 were not able to be recovered. One haplotype of chromosome 2 shares a transposable element with chromosome 3 and so is linked in the graph.

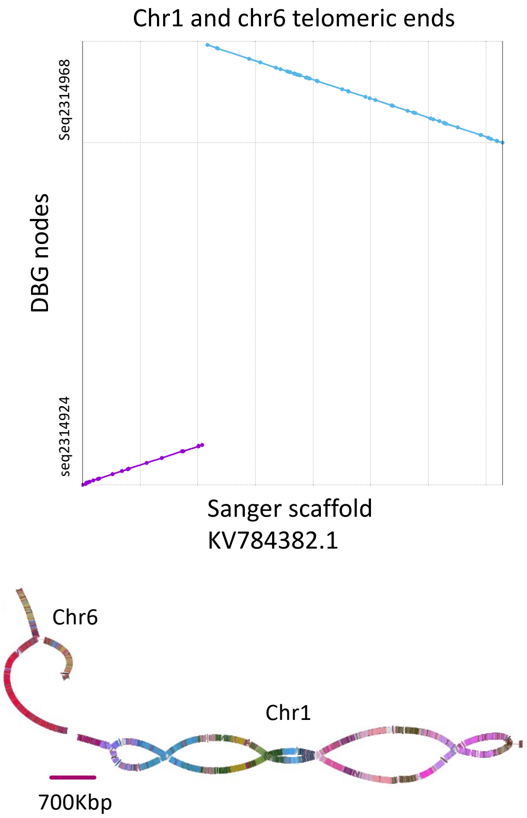


**Fig. S5.** Ends of the DBG chromosomes 1 and 6 aligned against a single scaffold from the published Sanger assembly. (Top) In the published assembly, the terminal end of chromosome 6 has been reverse complimented and joined through the telomere to chromosome 1. The remainder of chromosomes 1 and 6 in the published assemblies lie on different scaffolds. (Bottom) The DBG assembly karyotype is coloured by aligned contigs from the Sanger assembly, such that each contig is represented in a different colour. This also shows the previous level of assembly fragmentation in the Sanger assembly. The mis-assembly can be seen between the terminal ends of the chromosomes, which share the same colour.

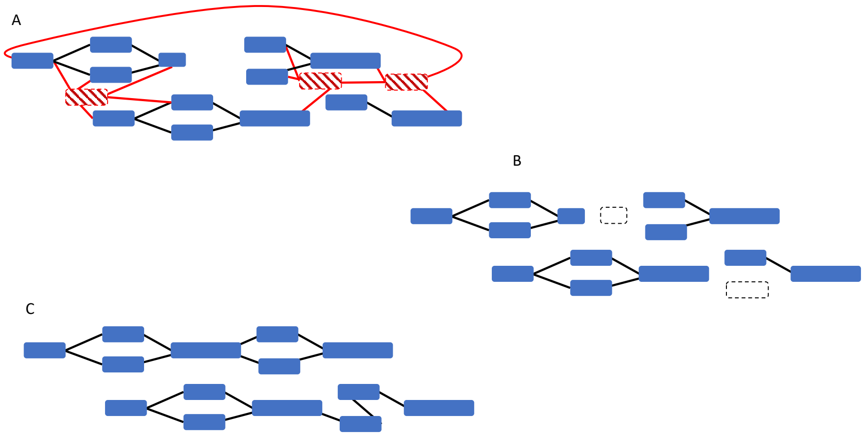


**Fig. S6.** Manual analysis of the De Bruijn Graph. A) The graph contains mis-joins and is difficult to follow. Red nodes are repeats, with a minimum of 2 input and 2 output links. As these are repeat regions, they can manually be deselected to simplify the graph. B) The graph is still fragmented because some nodes have been deselected due to not meeting the parameter requirements. This may be due to coverage, length or because they have been classified as repeats. These can be manually re-selected. C) After manual analysis, the graph is much simpler. The parametrisations can be changed to account for the manually adjusted nodes.

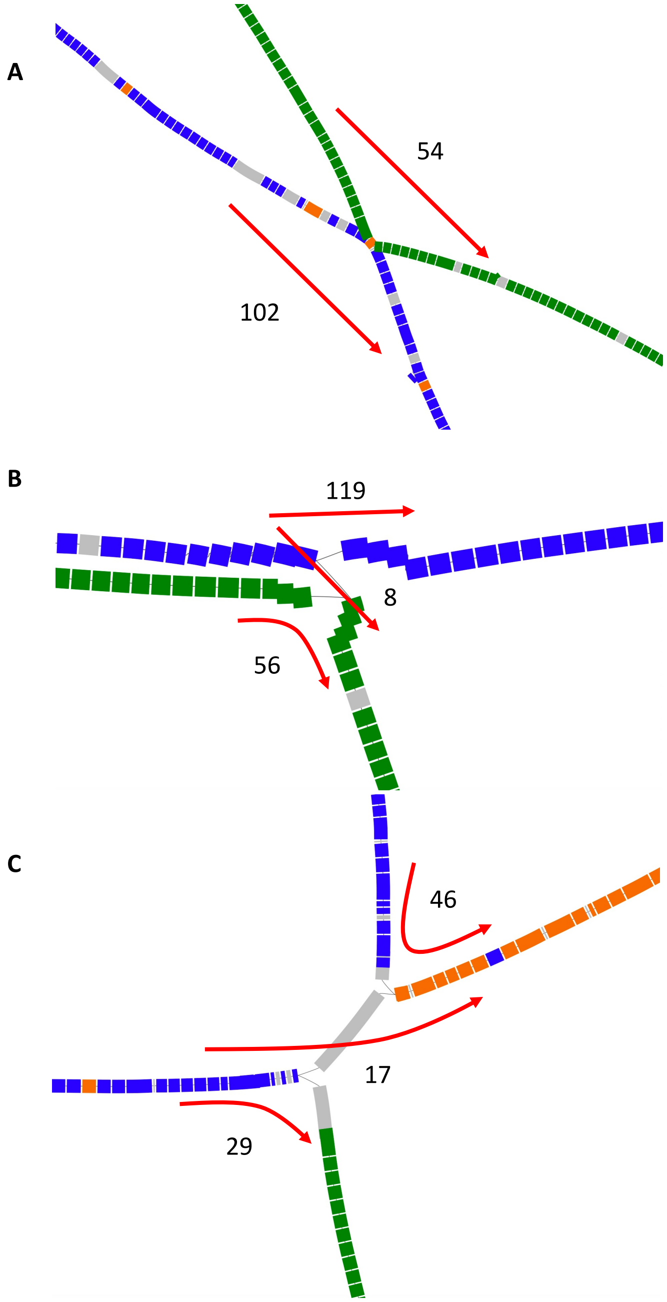


**Fig. S7.** Three types of crossovers identified in the assembly graph and the quantities of long reads that support each linkage. A) The 1x and 2x coverage regions converge into the 3x coverage node, before diverging out into 1x and 2x coverage regions again. Red arrows indicate the long-read support over the crossover with 54 reads supporting a 1x to 1x linkage and 102 reads supporting a 2x to 2x linkage. No long reads supported any other linkages. This is a small region of homozygosity between the bubbles. B) A mis-linked region determined not to be a genuine haplotypic crossover. Blue 2x coverage is supported by 119 long reads, and green 1x coverage is supported by 56, in roughly the quantities we would expect. An additional 7 long reads supported the unseen link 1x to 2x, which suggests this area is prone to sequencing errors. C) Complex crossover on chromosome 10. Two double-coverage bubble sides (blue) can be seen diverging into unique (green) or triple coverage (orange) regions. The quantities of long reads supporting these linkages suggest that there are four haplotypes here, three which converge into the same run of homozygosity, and one which diverges into a unique haplotype.
